# Comparative Genomics of *Vibrio vulnificus, Vibrio navarrensis*, and *Vibrio cidicii* Reveals Taxonomic Boundaries and Divergent Virulence Mechanisms

**DOI:** 10.1101/2025.07.25.666834

**Authors:** Keri Ann Lydon, Megan E. J. Lott

**Author notes:** 150 E Green St., Athens, GA, 30602.

## Abstract

Vibrionaceae are a diverse family of bacteria that contain pathogenic species, including those within the Vulnificus clade: *Vibrio vulnificus, Vibrio navarrensis*, and *Vibrio cidicii*. While *V. vulnificus* is a generally well characterized environmental pathogen, *V. cidicii* and *V. navarrensis* are relatively rare, recently identified species that our current understanding of virulence and environmental adaptation is limited. Here, we investigate genetic relatedness across these three species to identify shared and species-specific genes, including markers of virulence using publicly available genome assemblies. We evaluated phylogenetic and genomic diversity across this clade by sampling all available *V. navarrensis* and *V. cidicii* genomes, and a biodiverse curated set of four *V. vulnificus* ecotypes to ensure representative coverage. Our results indicate that all three species share 2,321 universally conserved genes, many of which are core bacterial functions. Moreover, *V. cidicii* and *V. navarrensis* have extensive genetic similarity between them, including average nucleotide identities >95% and 619 shared genes. Despite this similarity, they both remain more phylogenetically distant from *V. vulnificus* and lack key virulence genes such as *rtxA*, indicating alternative pathogenic mechanisms. Overall, these findings reveal that virulence potential varies across the clade and environmental adaptation potential varies between species and biotypes.

**IMPORTANCE:** *Vibrio* species are important environmental aquatic bacteria that pose a threat to human and animal health across the globe. This study applied comparative genomics to investigate the genetic relatedness of *Vibrio vulnificus, Vibrio navarrensis*, and *Vibrio cidicii*, with special focus on genes associated with environmental adaptation and virulence between and within each species. Results indicate *V. navarrensis* and *V. cidicii* share many genes and are phylogenetically close, and that they possess different virulence potential than *V. vulnificus*. This adds to our understanding of genetic diversity and pathogenic mechanisms within an important group of marine pathogens.

## INTRODUCTION

The family Vibrionaceae encompasses a wide range of nine genera made up of 51 clades and over 190 species, including many that are human and animal pathogens (Jiang et al., 2022). In the Vulnificus clade, *V. vulnificus* was originally classified as the sole species (Sawabe et al., 2007); however, recent advancements in microbial genomics have revealed newly described and closely related *Vibrio* species (Sawabe et al., 2013; Jiang et al., 2022), including *Vibrio navarrensis* (Urdaci et al., 1991) and *Vibrio cidicii* (Orata et al., 2016). Among these three species, *V. vulnificus* is clinically significant due to its severe pathogenic potential as one of the leading causes of death related to foodborne illness and responsible for 95% of all deaths from seafood consumption (Baker-Austin and Oliver, 2018). Additionally, *V. vulnificus* infections have a high case fatality rate (up to 50%) and result in severe outcomes such as necrotizing skin infections, sepsis, amputation, and death (Oliver, 2005). Beyond foodborne illness, many infections arise from direct contact with brackish waters and marine life (Jones and Oliver, 2009). Despite this well-characterized pathogenicity of *V. vulnificus*, the virulence potential and environmental ecology *V. cidicii* and *V. navarrensis* remain poorly understood, presenting a significant knowledge gap for public health risk assessment.

When compared to *V. vulnificus, V. navarrensis* has only recently been recognized as a human pathogen and is less frequently isolated, though it can be found in similar marine environments, sewage, and has been recovered from human blood, stool, and wound samples (Gladney et al., 2014; Gladney and Tarr, 2014; Schwartz et al., 2021). Beyond human illness, *V. navarrensis* has been linked to livestock (pigs and cattle) that suffered spontaneous abortions (Schwartz et al., 2017). In contrast, *V. cidicii* has primarily been isolated from aquatic environments and has not been classified as a human pathogen, though it has been isolated from human blood (Orata et al., 2016). Pathogenic potential across these species remains poorly characterized, despite recent clinical cases documented for *V. navarrensis* and *V. cidicii* (CDC, 2021).

*V. cidicii* is similar to *V. navarrensis* in most phenotypical tests, except for their ability to utilize L-Rhamnose as a sole carbon source (Orata et al., 2016). One recent study found *V. navarrensis* in high abundance across spatial and temporal gradients in surface waters, indicating diverse environment adaptation potential (Wan et al., 2025). Additionally, *V. navarrensis* has been found in biofilms on microplastics (Lai et al., 2022; Lo et al., 2025). Despite these environmental findings, our understanding of environmental drivers of abundance for these two species remains limited.

To better understand these species relationships and pathogenic potential, this study has three primary objects: 1) determine phylogenetic relatedness across the full spectrum of available *V. navarrensis* and *V. cidicii* genomes compared with a subset of diverse *V. vulnificus* strains, 2) identify species-specific genetic features that may explain differences in virulence potential and environmental adaptation, and 3) characterize the genomic diversity within each species to understand ecological and clinical strain clustering. We hypothesize that the three species within the Vulnificus clade represent distinct evolutionary lineages with species-specific adaptations for pathogenicity and environmental survival, despite their close phylogenetic relationship. This research provides a comprehensive assessment using comparative genomics to identify the genetic basis for observed differences in virulence potential and ecological distribution.

## MATERIALS AND METHODS

### Strain Selection

In this study, we obtained publicly available whole genome assemblies from the National Center for Biotechnology Information (NCBI) in September of 2024 for the three species that make up the Vulnificus clade: *Vibrio vulnificus, Vibrio navarrensis*, and *Vibrio cidicii* (Jiang et al., 2022). After the initial download, we discovered during curation that several of the isolates were duplicated, either through using unique identifiers created by the submitter or through multiple sequencing efforts used to build complete genomes. To limit the redundancy of genome sequences, we dereplicated the isolates by strain identification numbers to include only the complete final genome for each isolate, if one was available. If the NCBI quality control was flagged for not passing quality checks, we used the available draft genome (e.g. contig-level assembly) in its place. No isolates flagged for contamination or quality issues in NCBI were included in the study.

To ensure robust comparative analysis across species with vastly different representation in public databases, we employed a complete sampling strategy. We obtained all available genomes for the rarer species (*V. navarrensis* and *V. cidicii*) to eliminate sampling bias, while implementing curated biological diversity sampling for *V. vulnificus* to capture ecological and pathogenic diversity across established ecotypes. This approach prevents overrepresentation of *V. vulnificus* while maximizing genetic diversity representation across the clade. For *V. vulnificus*, we selected a subset of genomes (n=16) evenly across previously described ecotypes C1, C2, C3, and C4 (López-Pérez et al., 2019), making sure to include environmental and clinical isolates for each ecotype. In total, we obtained 62 isolates across the Vulnificus clade that included all isolates currently available for *V. navarrensis* (n=38) and *V. cidicii* (n=8). Accession numbers for genomes are listed in Table 1 with strain metadata and the original studies where the strains were previously characterized.

**Table 1.**
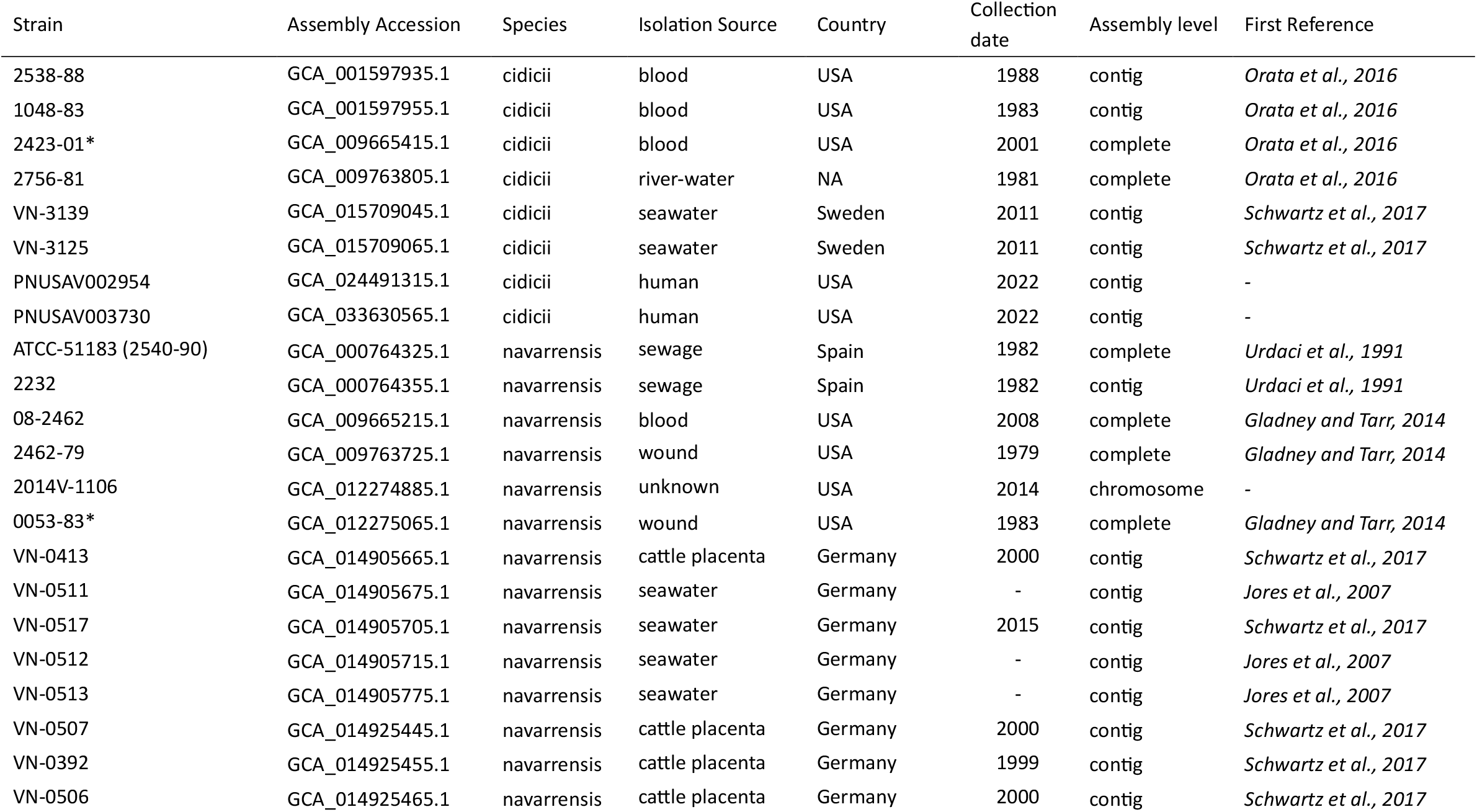
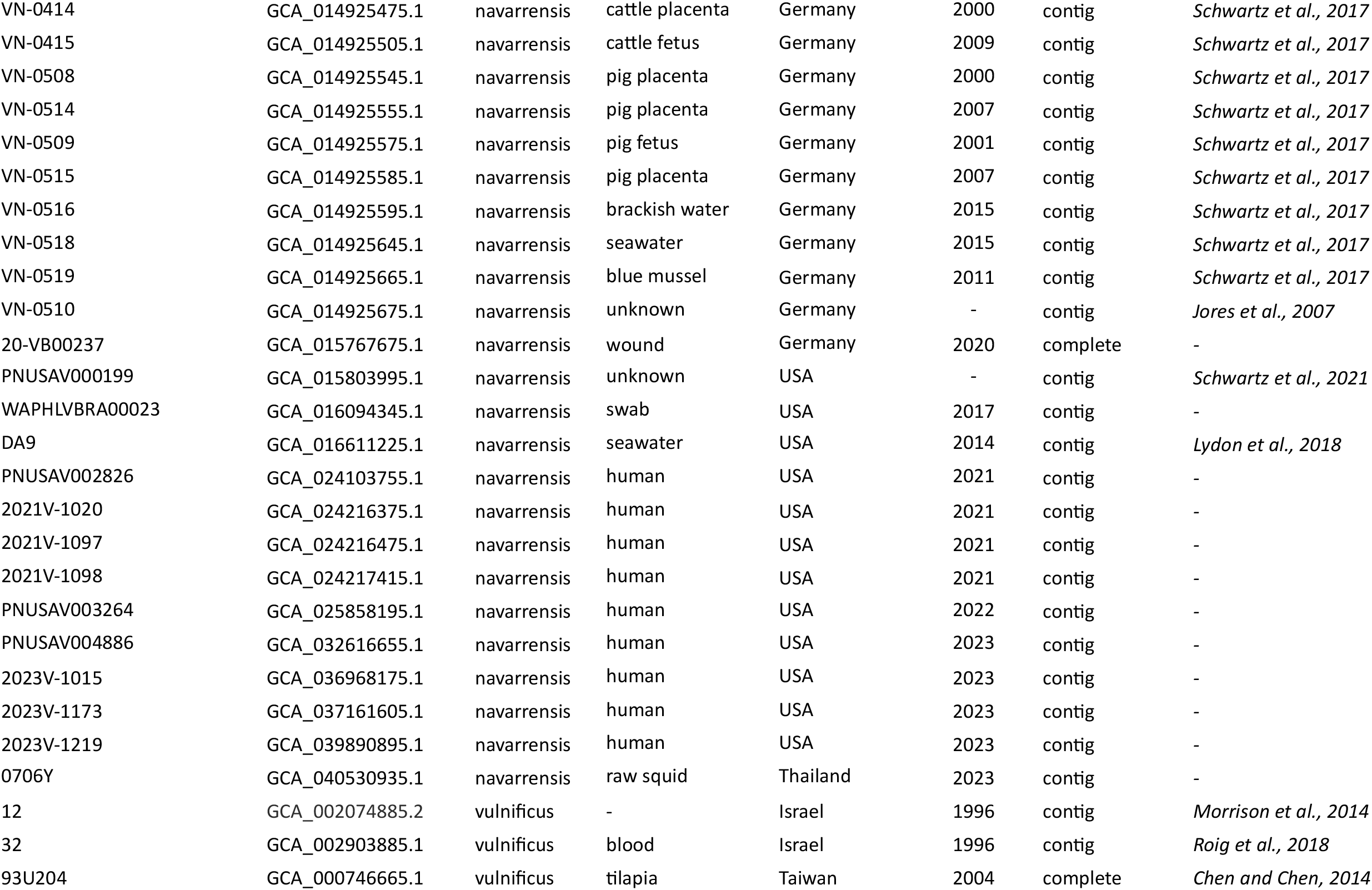
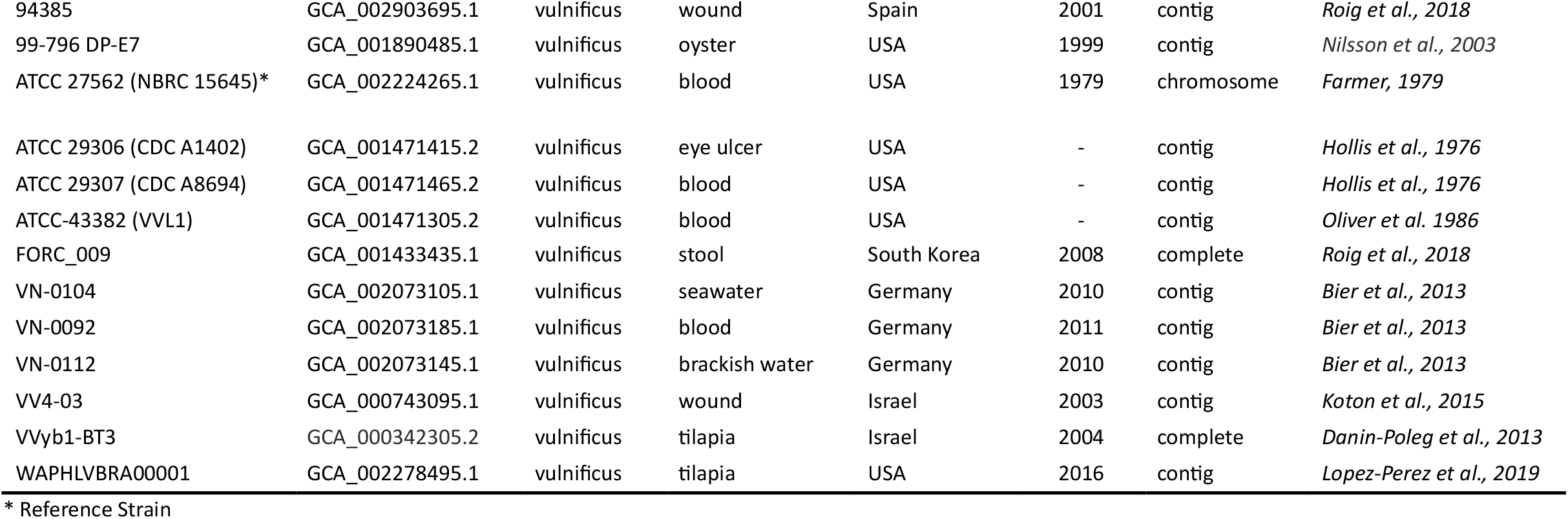
*Vibrio vulnificus, Vibrio navarrensis*, and *Vibrio cidicii* strains used in this study with the corresponding accession number for each assembly and the reference for each isolate’s initial appearance in the literature.

| Strain | Assembly Accession | Species | Isolation Source | Country | Collection date | Assembly level | First Reference |
| --- | --- | --- | --- | --- | --- | --- | --- |
| 2538-88 | GCA_001597935.1 | cidicii | blood | USA | 1988 | contig | *Orata et al., 2016* |
| 1048-83 | GCA_001597955.1 | cidicii | blood | USA | 1983 | contig | *Orata et al., 2016* |
| 2423-01* | GCA_009665415.1 | cidicii | blood | USA | 2001 | complete | *Orata et al., 2016* |
| 2756-81 | GCA_009763805.1 | cidicii | river-water | NA | 1981 | complete | *Orata et al., 2016* |
| VN-3139 | GCA_015709045.1 | cidicii | seawater | Sweden | 2011 | contig | *Schwartz et al., 2017* |
| VN-3125 | GCA_015709065.1 | cidicii | seawater | Sweden | 2011 | contig | *Schwartz et al., 2017* |
| PNUSAV002954 | GCA_024491315.1 | cidicii | human | USA | 2022 | contig | - |
| PNUSAV003730 | GCA_033630565.1 | cidicii | human | USA | 2022 | contig | - |
| ATCC-51183 (2540-90) | GCA_000764325.1 | navarrensis | sewage | Spain | 1982 | complete | *Urdaci et al., 1991* |
| 2232 | GCA_000764355.1 | navarrensis | sewage | Spain | 1982 | contig | *Urdaci et al., 1991* |
| 08-2462 | GCA_009665215.1 | navarrensis | blood | USA | 2008 | complete | *Gladney and Tarr, 2014* |
| 2462-79 | GCA_009763725.1 | navarrensis | wound | USA | 1979 | complete | *Gladney and Tarr, 2014* |
| 2014V-1106 | GCA_012274885.1 | navarrensis | unknown | USA | 2014 | chromosome | - |
| 0053-83* | GCA_012275065.1 | navarrensis | wound | USA | 1983 | complete | *Gladney and Tarr, 2014* |
| VN-0413 | GCA_014905665.1 | navarrensis | cattle placenta | Germany | 2000 | contig | *Schwartz et al., 2017* |
| VN-0511 | GCA_014905675.1 | navarrensis | seawater | Germany | - | contig | *Jores et al., 2007* |
| VN-0517 | GCA_014905705.1 | navarrensis | seawater | Germany | 2015 | contig | *Schwartz et al., 2017* |
| VN-0512 | GCA_014905715.1 | navarrensis | seawater | Germany | - | contig | *Jores et al., 2007* |
| VN-0513 | GCA_014905775.1 | navarrensis | seawater | Germany | - | contig | *Jores et al., 2007* |
| VN-0507 | GCA_014925445.1 | navarrensis | cattle placenta | Germany | 2000 | contig | *Schwartz et al., 2017* |
| VN-0392 | GCA_014925455.1 | navarrensis | cattle placenta | Germany | 1999 | contig | *Schwartz et al., 2017* |
| VN-0506 | GCA_014925465.1 | navarrensis | cattle placenta | Germany | 2000 | contig | *Schwartz et al., 2017* |
| VN-0414 | GCA_014925475.1 | navarrensis | cattle placenta | Germany | 2000 | contig | *Schwartz et al., 2017* |
| VN-0415 | GCA_014925505.1 | navarrensis | cattle fetus | Germany | 2009 | contig | *Schwartz et al., 2017* |
| VN-0508 | GCA_014925545.1 | navarrensis | pig placenta | Germany | 2000 | contig | *Schwartz et al., 2017* |
| VN-0514 | GCA_014925555.1 | navarrensis | pig placenta | Germany | 2007 | contig | *Schwartz et al., 2017* |
| VN-0509 | GCA_014925575.1 | navarrensis | pig fetus | Germany | 2001 | contig | *Schwartz et al., 2017* |
| VN-0515 | GCA_014925585.1 | navarrensis | pig placenta | Germany | 2007 | contig | *Schwartz et al., 2017* |
| VN-0516 | GCA_014925595.1 | navarrensis | brackish water | Germany | 2015 | contig | *Schwartz et al., 2017* |
| VN-0518 | GCA_014925645.1 | navarrensis | seawater | Germany | 2015 | contig | *Schwartz et al., 2017* |
| VN-0519 | GCA_014925665.1 | navarrensis | blue mussel | Germany | 2011 | contig | *Schwartz et al., 2017* |
| VN-0510 | GCA_014925675.1 | navarrensis | unknown | Germany | - | contig | *Jores et al., 2007* |
| 20-VB00237 | GCA_015767675.1 | navarrensis | wound | Germany | 2020 | complete | - |
| PNUSAV000199 | GCA_015803995.1 | navarrensis | unknown | USA | - | contig | *Schwartz et al., 2021* |
| WAPHLVBRA00023 | GCA_016094345.1 | navarrensis | swab | USA | 2017 | contig | - |
| DA9 | GCA_016611225.1 | navarrensis | seawater | USA | 2014 | contig | *Lydon et al., 2018* |
| PNUSAV002826 | GCA_024103755.1 | navarrensis | human | USA | 2021 | contig | - |
| 2021V-1020 | GCA_024216375.1 | navarrensis | human | USA | 2021 | contig | - |
| 2021V-1097 | GCA_024216475.1 | navarrensis | human | USA | 2021 | contig | - |
| 2021V-1098 | GCA_024217415.1 | navarrensis | human | USA | 2021 | contig | - |
| PNUSAV003264 | GCA_025858195.1 | navarrensis | human | USA | 2022 | contig | - |
| PNUSAV004886 | GCA_032616655.1 | navarrensis | human | USA | 2023 | contig | - |
| 2023V-1015 | GCA_036968175.1 | navarrensis | human | USA | 2023 | contig | - |
| 2023V-1173 | GCA_037161605.1 | navarrensis | human | USA | 2023 | contig | - |
| 2023V-1219 | GCA_039890895.1 | navarrensis | human | USA | 2023 | contig | - |
| 0706Y | GCA_040530935.1 | navarrensis | raw squid | Thailand | 2023 | contig | - |
| 12 | GCA_002074885.2 | vulnificus | - | Israel | 1996 | contig | *Morrison et al., 2014* |
| 32 | GCA_002903885.1 | vulnificus | blood | Israel | 1996 | contig | *Roig et al., 2018* |
| 93U204 | GCA_000746665.1 | vulnificus | tilapia | Taiwan | 2004 | complete | *Chen and Chen, 2014* |
| 94385 | GCA_002903695.1 | vulnificus | wound | Spain | 2001 | contig | *Roig et al., 2018* |
| 99-796 DP-E7 | GCA_001890485.1 | vulnificus | oyster | USA | 1999 | contig | *Nilsson et al., 2003* |
| ATCC 27562 (NBRC 15645)* | GCA_002224265.1 | vulnificus | blood | USA | 1979 | chromosome | *Farmer, 1979* |
| ATCC 29306 (CDC A1402) | GCA_001471415.2 | vulnificus | eye ulcer | USA | - | contig | *Hollis et al., 1976* |
| ATCC 29307 (CDC A8694) | GCA_001471465.2 | vulnificus | blood | USA | - | contig | *Hollis et al., 1976* |
| ATCC-43382 (VVL1) | GCA_001471305.2 | vulnificus | blood | USA | - | contig | *Oliver et al. 1986* |
| FORC_009 | GCA_001433435.1 | vulnificus | stool | South Korea | 2008 | complete | *Roig et al., 2018* |
| VN-0104 | GCA_002073105.1 | vulnificus | seawater | Germany | 2010 | contig | *Bier et al., 2013* |
| VN-0092 | GCA_002073185.1 | vulnificus | blood | Germany | 2011 | contig | *Bier et al., 2013* |
| VN-0112 | GCA_002073145.1 | vulnificus | brackish water | Germany | 2010 | contig | *Bier et al., 2013* |
| VV4-03 | GCA_000743095.1 | vulnificus | wound | Israel | 2003 | contig | *Koton et al., 2015* |
| VVyb1-BT3 | GCA_000342305.2 | vulnificus | tilapia | Israel | 2004 | complete | *Danin-Poleg et al., 2013* |
| WAPHLVBRA00001 | GCA_002278495.1 | vulnificus | tilapia | USA | 2016 | contig | *Lopez-Perez et al., 2019* |

\* Reference Strain

### Genome Analyses

All genome assemblies were processed with Galaxy (Afgan, et al., 2018; The Galaxy Community, 2024) on the public server UseGalaxy.eu; unless noted, default parameters were used. We evaluated genomic features and assembly quality with QUAST v5.3.0 (Gurevich et al., 2013; Mikeenho et al., 2016; Mikheenko, et al., 2018) and used Prokka v1.14.6 (Cuccuru et al., 2014; Seemann, 2014) to annotate genomes for downstream analyses. High quality genome assemblies were defined as those with N50 values >50,000 bp and <301 contigs, assemblies meeting these criteria were included in subsequent analyses.

To identify conserved orthologous genes across the Vulnificus clade, we processed all genome annotations with ROARY v3.13.0 (Page et al., 2015). ROARY was specifically chosen for its conservative gene clustering approach, which has been successfully used for interspecies comparative analyses within bacterial genera (Smet et al., 2018). We adjusted ROARY’s default parameters to ensure better detection of orthologous groups for interspecies comparisons by reducing the identity threshold to 70% (Donati et al., 2010) and MCL inflation to 1.4 to reduce the number of false positives. Core genes were defined as those present in all three species (99% of isolates), and paralogs were not split to focus on functional gene presence rather than copy number variation.

We used IQ-TREE v2.4.0 (Minh et al., 2020) with ModelFinder (Kalyaanamoorthy et al., 2017) and 1000 ultrafast bootstraps (Hoang et al., 2017) to infer the best-scoring maximum likelihood tree from the core gene alignment of universally conserved genes produced with ROARY. We rooted the tree at the midpoint and visualized it with iTOL v7 (Letunic and Bork, 2024). Average nucleotide identity (ANI) was estimated using fastANI v1.3 (Jain et al., 2018), then visualized as a heatmap with R v. 4.5.0, Rstudio (2025.05.1+513), using packages ‘ggdendro’ and ‘ggplot’. The heatmap was clustered by inferring a dendrogram based on hierarchical clustering (de Vries and Ripley 2024) of the gene presence-absence table produced by ROARY.

To determine which genes were significantly associated with each species and isolation sources (clinical, water, aquatic animal, livestock), we used SCOARY v1.6.16 (Tang et al., 2020). Briefly, each species or category was depicted as a discrete binomial trait (i.e., belonging to the species or not). Genes were reported if they were 100% specific and had a p-value <0.05 for naive-p, Bonferroni, and Benjamin-Hochberg tests. Significant genes with high specificity were then summarized into biologically meaningful groups by species.

For functional classification, the core gene alignment was processed into a single consensus sequence using the ‘Consensus sequence from aligned FASTA’ tool v.1.0.0 (Keck, 2020), with the most frequent nucleotide as the consensus model. The consensus sequence was then processed with eggNOG v. 2.1.8 (Huerta-Cepas et al., 2016; Huerta-Cepas et al., 2017) to determine KEGG orthology (Kanehisa, 2016) and gene ontology (GO) categories (Ashburner et al., 2000). Clusters of Orthologous Groups (COG) categories for universally conserved genes were visualized in R (R Core Team 2021).

### Virulence Factors and Antimicrobial Resistance

We screened each strain individually with ABRicate v1.0.1 (Seeman, 2016) for the presence of virulence factors using the Virulence Factors of Pathogenic Bacteria (VFDB) (Chen et al., 2005) database and antimicrobial resistance genes against the Comprehensive Antibiotic Resistance Database (CARD) (Jia et al., 2017), with 70% minimum percent identity and coverage.

### Data Availability

All accession numbers for genome assemblies used in this study can be found in Table 1. Supplemental materials and datafiles are available at OSF: https://osf.io/bh8y2/?view_only=946c3c2250114a16ba02f34baab7a115.

## RESULTS

### Genomic Features

In this study, we evaluated the relationship between species within the Vulnificus clade, including *V. vulnificus, V. navarrensis*, and *V. cidicii*. Assembly quality metrics (e.g., contig counts, N50 values, and total assembly lengths) are provided in Supplementary Table 1. Examination of genomic features indicated that Vulnificus clade genomes (n=62) were on average 4,700,000 bp (range 4,139,000 - 5,745,000 bp), with a mean GC content of 47.8% (range 46.4 - 48.6%). When looking at each species individually, *V. vulnificus* (n=16) had the largest mean genome size of 5,059,000 bp (range 4,680,000 – 5,745,000 bp), while *V. navarrensis* (n=38) and *V. cidicii* (n=8) were 4,564,000 bp (range 4,139,000 – 5,110,000 bp) and 4,623,000 bp (range 4,390,000 – 5,048,000 bp), respectively. Mean GC content was 46.7% (range 46.4 - 47.9%) for *V. vulnificus* isolates, and 48.3% (range 47.7 - 48.6%) and 48.0% (range 46.7 - 48.3%) each for *V. navarrensis* and *V. cidicii*, respectively.

### Phylogenetic Analysis

ModelFinder determined the best-fit model for maximum likelihood phylogeny reconstruction across 1,173,000 nucleotide sites from universally conserved genes to be GTR+F+I+R6. The resulting phylogenetic tree revealed clear interspecies relationships with well-supported clades (Fig. 1). The tree exhibited a clear bifurcating structure with *V. vulnificus* forming one monophyletic clade, with all 4 ecotype clusters resolved, and *V. navarrensis* and *V. cidicii* forming a separate sister clade. All species-level nodes achieved 100% bootstrap support values.

**Figure 1.**
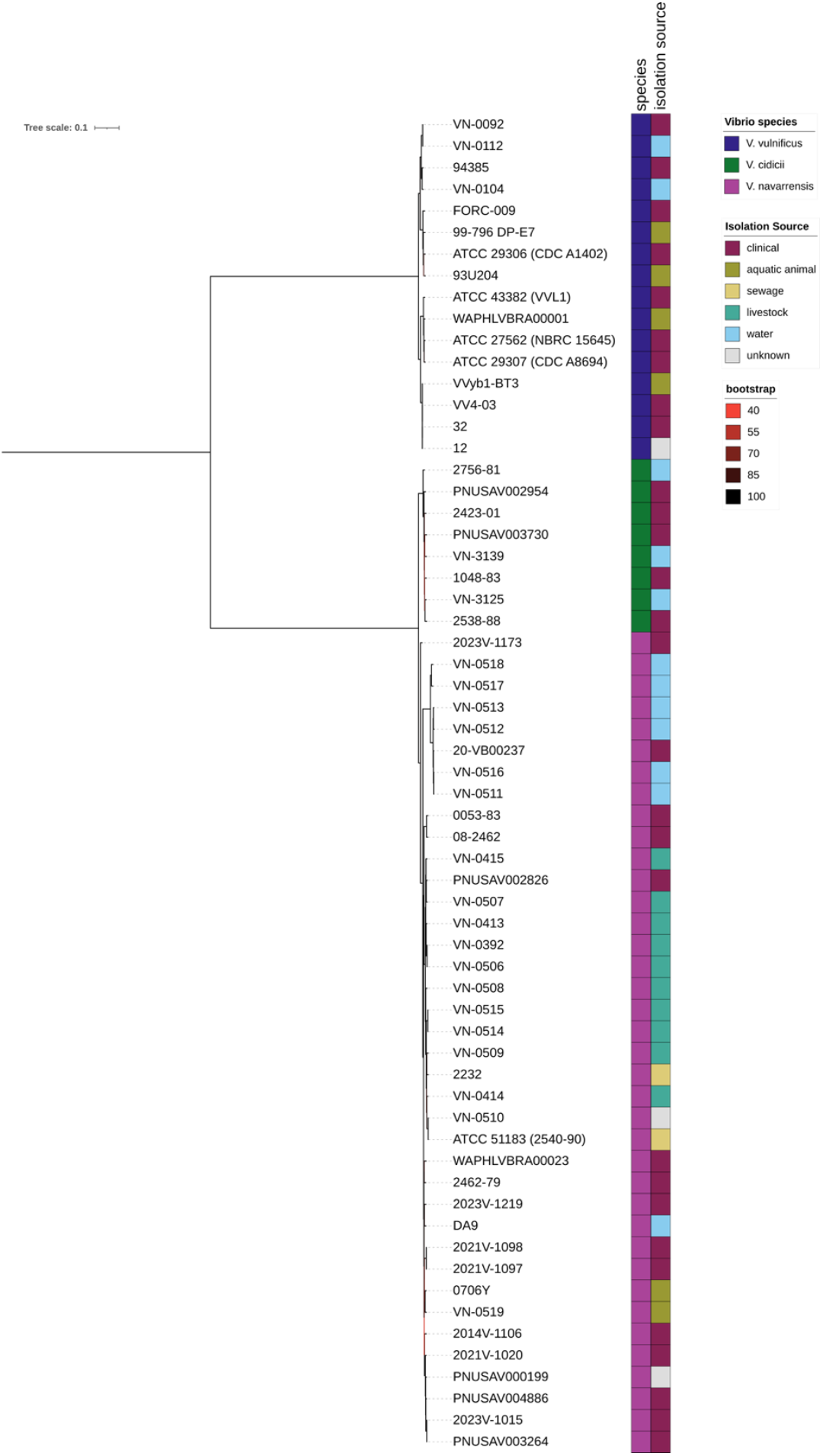
Phylogenetic tree inferred from the concatenated alignment of universally conserved genes for all three species of the Vulnificus clade: *Vibrio vulnificus, Vibrio navarrensis*, and *Vibrio cidicii*. Maximum likelihood phylogeny was constructed using IQ-TREE v2.4.0 with ModelFinder (GTR+F+I+R6 model) and 1,000 ultrafast bootstraps from 1,173,000 nucleotide sites. The tree is midpoint-rooted. Strain names and isolation sources are indicated for each isolate.

*V. navarrensis* displayed high intraspecies genetic diversity, forming multiple phylogenetic subclades with bootstrap values ranging from 47 to 100%. Lower bootstrap values were concentrated at nodes within the *V. navarrensis* clade, particularly among environmental and livestock-associated lineages. Livestock-associated isolates from cattle and pig spontaneous abortion cases showed phylogenetic clustering. In contrast, environmental isolates from seawater, sewage, and brackish water were phylogenetically dispersed throughout the tree. Clinical isolates from blood, wound, stool, and unspecified clinical sources were distributed across multiple phylogenetic lineages. Of particular interest, a recent clinical isolate (PNUSAV002826, isolated in 2021) clustered with livestock isolates from cattle fetal and placental tissues. Similarly, environmental isolate *V. navarrensis* DA9 from a coastal canal in the Florida Keys (USA) clustered closely with recent USA clinical isolates 2021V-1098, 2021V-1097, and 2023V-1219.

In contrast, hierarchical clustering based on gene presence/absence at 70% identity threshold showed clusters of *V. navarrensis* and *V. cidicii* isolates intermixed (Fig. 2), contrasting with the clear phylogenetic separation observed in the core genome phylogenetic tree.

**Figure 2.**
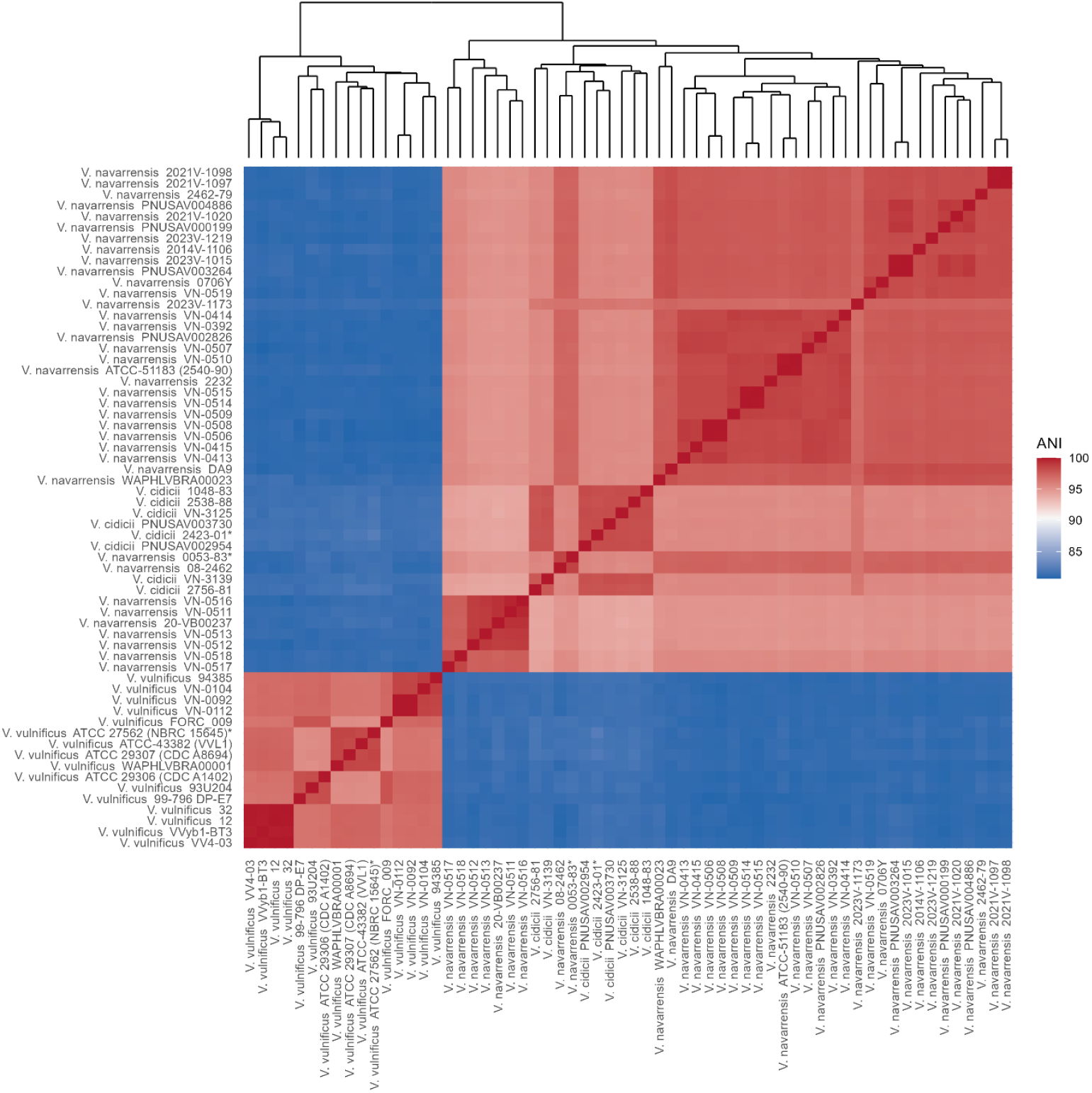
Pairwise average nucleotide identity (ANI) across the Vulnificus clade. ANI values were calculated using fastANI v1.3 and visualized as a heatmap with hierarchical clustering. The dendrogram indicates clustering based on gene presence-absence profiles produced by ROARY at 70% identity threshold.

### Average Nucleotide Identity (ANI) Analysis

Across the Vulnificus clade, Average Nucleotide Identity (ANI) was on average 90.8% with a range of 80.7% to 100% (Fig. 2). Intraspecies comparisons revealed higher ANI values, as expected. We observed average intraspecies ANI values of 98.1% (SD = 0.1), 97.1% (SD = 1.4), and 97.0% for *V. cidicii, V. navarrensis*, and *V. vulnificus*, respectively. Interspecies pairwise comparisons identified distinct patterns; both *V. cidicii* and *V. navarrensis* had lower ANI values when compared to *V. vulnificus* (81.6% ± 0.2% and 81.1% ± 0.2%, respectively). Contrarily, the pairwise comparison between *V. navarrensis* and *V. cidicii* had an average ANI value of 95.3% (SD = 0.5). Within *V. navarrensis*, biotype *pommerensis* strains showed almost identical average ANI values of 95.1% (SD = 0.2) when compared to all non-pommerensis *V. navarrensis* isolates. *V. navarrensis* biotype *pommerensis* strains among themselves had average ANI values of 98.5% (SD = 1.0).

### Universally Conserved Genes

We identified 2,321 universally conserved genes (99 - 100% of strains) across all three species. An additional 189 highly conserved genes (95 - 99% of strains), 2,767 shell genes (15 - 95% of strains), and 9,358 cloud genes (0 - 15% of strains) were found across all isolates. Functional annotation of universally conserved genes with eggNOG-mapper assigned COG categories to 1,267 genes, representing all major COG functional categories (Fig. 3). Metabolism-related functions were the largest category (38%), followed by cellular processes and signaling (23.6%), and information storage and processing (20.9%). The largest singular COG categories were those that were poorly characterized with unknown functions (S). This was followed by core bacterial processes of transcription (K), translation, ribosomal structure and biogenesis (J), and energy production and conversion (C). Interestingly, there were some conserved functions across Inorganic ion transport and metabolism (P).

**Figure 3.**
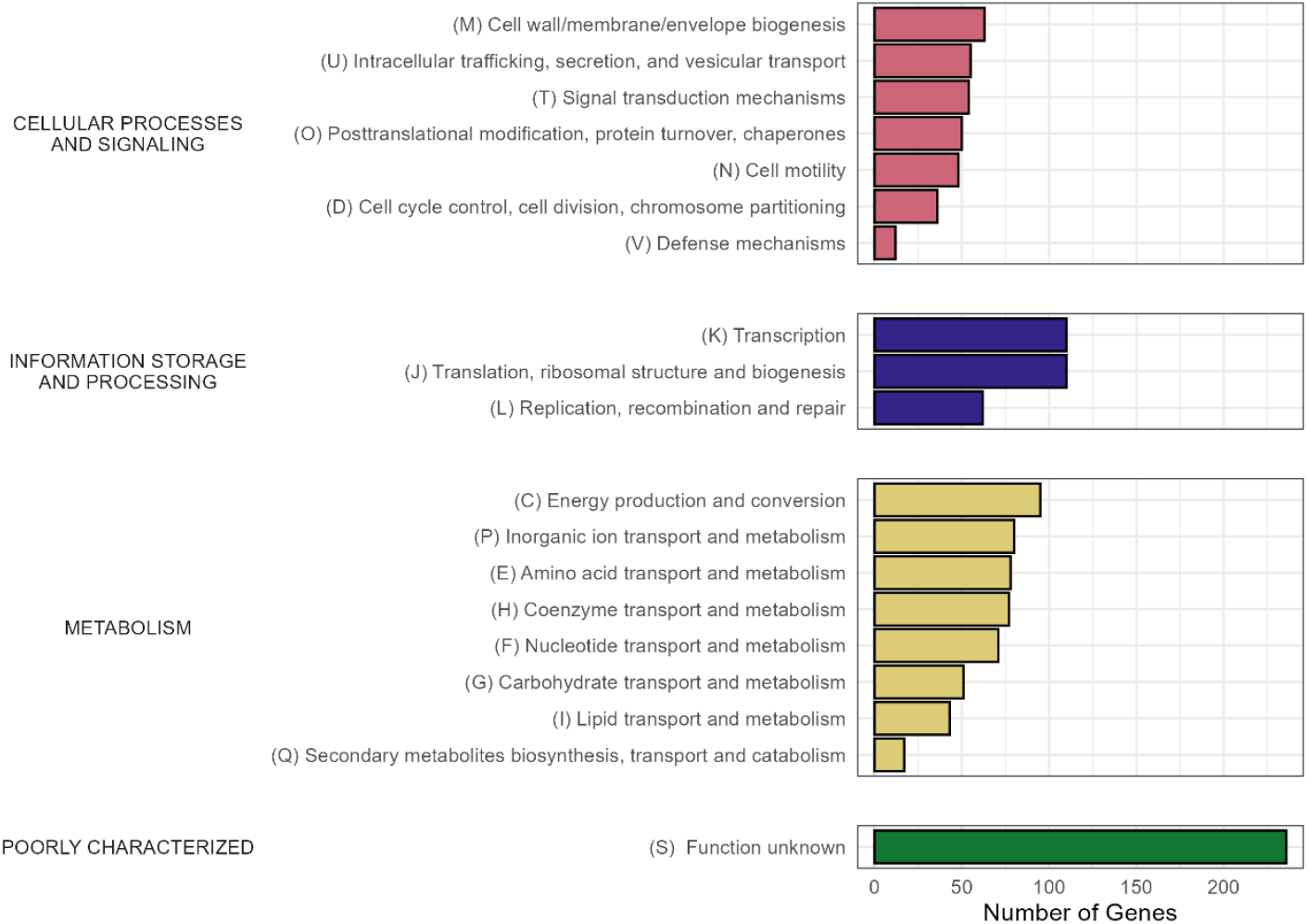
Functional categorization of universally conserved genes across the Vulnificus clade. The 2,321 universally conserved genes (present in 99-100% of strains) were functionally annotated using eggNOG-mapper v2.1.8 and assigned to COG (Clusters of Orthologous Groups) categories. Bars indicate the numbers of genes assigned to COG categories..

We identified many pathogenicity-associated genetic elements conserved across all three species. Complete flagellar biosynthesis and motility systems were present, including structural (*fliG, fliM, fliN, flgB, flgC, flgF, flgH, flgI*), regulatory (*fliA, fliS*), and chemotactic components (*cheA, cheY, cheZ*), alongside Type IV pilus assembly machinery (*pilB, pilD, pilU*). Intracellular communication capabilities included quorum sensing regulatory circuits (*luxS, luxR*, and *luxT*) and two-component regulatory systems (*cpxA, cpxR, phoQ, phoR*). Environmental adaptation mechanisms were equally well-conserved, including oxidative stress defense systems (*katG, sodB, oxyR*), iron acquisition pathways (*feoA, feoB, bfr*), and heavy metal resistance genes (*copA, zntA, znuB, znuC, zur*).

### Species Specific Features

We identified genes significantly associated with each species with SCOARY (Supplementary Table 2). For *V. vulnificus*, there were 1,020 genes specific to the species, including 720 hypothetical proteins. Among the 300 annotated genes, the largest biologically meaningful categories were nutrient acquisition and transport (47 genes), environmental stress response (26 genes), and regulation/signaling (35 genes). Notably, *V. vulnificus* was the only species to possess some virulence-associated genes, including cytolytic toxins (*ltxA, ltxB, ltxC*), RTX toxin (*rtxA*), hemolysin (*tdh_2*), and other virulence factors (*bepA* family).

We identified 619 genes shared between *V. cidicii* and *V. navarrensis*, of which 216 were hypothetical proteins. The 403 annotated shared genes represented some core functions known to *Vibrio* species, including iron acquisition, basic stress response, energy metabolism, and chemotaxis systems. For *V. cidicii* strains, 56 genes were specific to the species of which 39 were hypothetical proteins. *V. cidicii* showed metabolic specialization among its 17 annotated species-specific genes, particularly rhamnose catabolism (*rhaT, rha D, rhaB, rhaA, rhaM, rhaS_5*) and alcohol metabolism (*adhB_1*), as well as tetrathionate respiration capability (*ttrR, ttrS*). *V. navarrensis* had 49 genes that were species specific, with 26 hypothetical proteins. *V. navarrensis* had 23 annotated species-specific genes distributed across nutrient acquisition and environmental stress response categories, with no unique metabolic specializations identified.

There were no highly specific significant genes when we examined genomes by trait (clinical, water, aquatic animal, livestock).

### Antimicrobial resistance and virulence gene distribution

ABRicate screening revealed distinct antimicrobial resistance and virulence factor profiles. Screening against the CARD database revealed widespread resistance genes across all three species, with all strains carrying multiple resistance genes ranging from 2-5 genes per genome. Universal resistance genes included the cAMP receptor protein gene (*CRP*) and tetracycline resistance gene (*tet(35)*), present in all 62 strains. Chloramphenicol resistance (*catB9*) showed species-specific distribution, with high prevalence in *V. vulnificus* (15/16 strains) compared to *V. navarrensis* (2/38 strains) and *V. cidicii* (1/8 strains). Additional resistance genes included quinolone resistance (*qnrVC6*) in 3/62 strains and sulfamethoxazole resistance (*sul2*) in 1/62 strains. *Vibrio cholerae varG* demonstrated complete *V. vulnificus* specificity (16/16 strains). Conversely, the *ugd* gene, linked to polymyxin resistance, showed an inverse distribution pattern with higher prevalence in *V. navarrensis* (21/38 strains) compared to *V. vulnificus* (2/16 strains) and *V. cidicii* (3/8 strains).

VFDB analysis revealed species-specific differences in virulence gene distribution. While core Vulnificus clade virulence factors including immunolipoprotein A (*ilpA*), outer membrane protein U (*ompU*), and thermolabile hemolysin (*tlh*) were present across all three species, additional virulence genes showed clear species boundaries. *V. vulnificus* strains consistently carried more virulence genes (range: 14-29) compared to both *V. navarrensis* (range: 8-11) and *V. cidicii* (range: 8-11). Type VI Secretion System (T6SS) components were rarely detected across all three species, with complete gene clusters present in only 1/62 strains overall.

The most striking species difference was the complete restriction of RTX cytotoxin genes (*rtxB, rtxC, rtxD*) to *V. vulnificus*. All 16 *V. vulnificus* strains carried these genes while they were completely absent from all 38 *V. navarrensis* and 8 *V. cidicii* strains. This species-specific distribution was consistent across all ecological sources and geographic origins represented in the dataset. Similarly, the type IV pilus assembly gene (*pilT*) showed identical species restriction (16/16 *V. vulnificus*, 0/46 other species).

## DISCUSSION

### Phylogenetic Relationships and Taxonomic Revision

Our interspecies comparison across the Vulnificus clade provides insights into evolutionary relationships and functional divergence within this clinically important Vibrionaceae clade. We employed complete sampling of all available genomes for the rare species *V. navarrensis* and *V. cidicii*, and biologically diverse sampling for the well-characterized pathogen *V. vulnificus*. Previous work on phylogenetic relationships across this clade has been limited by the rare isolation of *V. navarrensis* and *V. cidicii*. All phylogenetic relationships compare to previous work (Jiang et al., 2022), however there has been continued emphasis on a close relationship between *V. navarrensis* and *V. cidicii* to *V. vulnificus* (Gladney and Tarr, 2014, Orata et al., 2016) without defining features that characterize that relationship.

In congruence with previous research (Hounmanou et al., 2025), our expanded genomic analysis with additional isolates confirms *V. vulnificus* is more evolutionarily distant from both *V. navarrensis* and *V. cidicii* than either of these two species are to each other. Furthermore, we have identified that many of the shared gene pathways within universally conserved genes across all three species represent core bacterial functions, though many remain uncharacterized. In contrast, *V. navarrensis* and *V. cidicii* contain a substantial number of shared genes (619) between them that are not present in *V. vulnificus*. These findings suggest a more complex evolutionary history, including a likely recent common ancestor and divergence of *V. cidicii* and *V. navarrensis*.

We have also identified that *V. cidicii* and *V. navarrensis* have high pairwise ANI values (>95%), close to the 95% threshold used for species delineation (Jain et al., 2018; Rodriguez-R et al., 2024) and species quality control at NCBI (Kannan et al., 2023). The average ANI values observed between *V. navarrensis* and *V. cidicii* (95.3%) and *V. navarrensis* biotype *pommerensis* and non-*pommerensis V. navarrensis* (95.1%) place both lineages in the transitional zone where species boundaries are typically established (Rodriguez-R et al., 2024), suggesting parallel evolutionary processes and recent speciation. This in combination with mixed hierarchical clustering of gene presence/absence indicates likely variable evolutionary pressures on the universally conserved genes and accessory genes in the sister clade. This evolutionary dynamic is common in aquatic bacteria and may reflect adaptation to overlapping but distinct ecological niches (Hunt et al., 2008). Future taxonomic assessments should consider both genomic and ecological evidence, particularly as additional isolates become available.

### Environmental Adaptations and Transmission Potential

*V. vulnificus* is notorious for not only being a foodborne pathogen, but also one that can infect via non-foodborne exposure pathways, including direct contact with marine animals and brackish waters (Jones and Oliver, 2009). *V. navarrensis* and *V. cidicii* have been isolated from human clinical samples, but less is known about the transmission pathways for these two pathogens. Our genomic findings, paired with previous environmental observations, suggest an aquatic pathway for exposure.

Globally, *V. navarrensis* has been identified in aquatic environments such as the Baltic Sea (Hounmanou et al., 2025), coastal areas near Hong Kong (Lo et al., 2025), the coastal Florida Keys (Lydon et al., 2018; Lydon et al., 2024), and from brackish water (Macián et al., 2000; Jores et al., 2007; Schwartz et al., 2017). Like *V. vulnificus*, human clinical isolates of *V. navarrensis* cluster with isolates collected from water samples, suggesting a waterborne transmission pathway. However, unlike *V. vulnificus, V. navarrensis* can grow at low salinities and without NaCl (Urdaci et al., 1991), indicating that exposures expand to include freshwater. *V. navarrensis* has been found in high abundance across the year in coastal areas where salinities ranged from 4.6-33.5 ppt (Wan et al., 2025). Conversely, *V. vulnificus* thrives from 5 ppt to 25 ppt salinity and is most often associated with brackish waters (Kaspar and Tamplin, 1993).

With very few isolates in existence, characterizing environmental patterns of *V. cidicii* reservoirs is limited. However, we can make some inferences on potential environmental adaptations from genetic evidence in this study. The combination of rhamnose catabolic genes and tetrathionate respiration genes as species-specific features of *V. cidicii* indicates they have metabolic capabilities for specialization in environments where rhamnose is present, such as algal exudates, bacterial exopolysaccharides, and dissolved organic matter (DOM) (Alderkamp et al., 2007; Mühlenbruch et al., 2018). Our results are supported by previous characterizations of *V. cidicii* being able to use rhamnose as a sole carbon source (Orata et al., 2016). This contrasts with both *V. vulnificus* and *V. navarrensis*, who did not share that characteristic.

### One Health Significance

These findings highlight the importance of a One Health approach to understanding environmental reservoirs and transmission dynamics. *V. vulnificus* is considered a zoonotic pathogen; the bacterium can be transmitted through dermal contact with or consumption of contaminated aquaculture products (Austin 2010, Hernandez-Cabanyero and Amaro 2022). The zoonotic potential of *V. navarrensis and V. cidicii* is not well understood. However, we found that two recent human clinical infections of *V. navarrensis* (*V. navarrensis* 2023V-1173 and *V. navarrensis* PNUSAV002826) clustered with livestock abortion cases when previously none had clustered in this way (Schwartz et al., 2021). These findings warrant further investigation into the transmission pathways of this pathogen.

Of note, we identified that most of the clinical isolates from the last decade in the USA cluster with *V. navarrensis* DA-9, an environmental isolate originating from seawater (Lydon et al., 2018; Lydon et al., 2024). There is also the possibility of sewage infrastructure influencing environmental abundance of *V. navarrensis;* phylogenetic clustering in this study is congruent with previous research where sewage and veterinary isolates were separate from water isolates (Schwartz et al., 2017).

### Virulence Mechanisms and Antimicrobial Resistance

Our comparative genome analysis identified several virulence-associated genes conserved, including iron acquisition systems and outer membrane proteins, which are often associated with host adaptations and immune evasion. Furthermore, all species in the clade were positive for hemolysin genes. However, RTX cytotoxin genes were restricted to *V. vulnificus*, with *rtxB/C/D* present in all *V. vulnificus* strains and *rtxA* absent in *V. navarrensis* and *V. cidicii*. This finding suggests a baseline of virulence potential shared across the clade associated with the host, but specialized cytotoxic virulence limited to *V. vulnificus*. It is likely that *V. navarrensis* and *V. cidicii* utilize the core functions to evade the host immune response, but beyond that there is no evidence of additional virulent traits. Our finding of complete RTX gene absence in *V. navarrensis* and *V. cidicii* contrasts with Schwartz et al., (2017), who reported RTX-positive *V. navarrensis* strains. This discrepancy likely stems from methodological differences and scale of analysis. Schwartz et al. used targeted BLASTN searches on a smaller subset of strains, while our study employed comprehensive screening across all available isolates using two independent approaches, ABRicate against the VFDB database and ROARY/SCOARY comparative analysis. Our broader sampling and dual method validation provides stronger evidence for the species-specific restriction of RTX toxins to *V. vulnificus*. Additionally, the use of stringent statistical criteria in our analysis reduces the likelihood of false-positive associations.

Antimicrobial resistance (AMR) genes were observed across the Vulnificus clade, with many genes shared across all species. The presence of AMR indicates that there is likely vertical transfer of some conserved genes from a common ancestor not tied to the isolation location. *V. vulnificus* has a wide range of AMR genes in terms of total number, which reflects interaction with humans and human infections (exposure to more antimicrobials).

### Study Limitations

Our study has some limitations inherent to comparative genomic analysis of publicly available data. Assembly quality varies across the 62 genomes, with a mixture of complete and draft assemblies potentially affecting gene presence/absence detection. However, our conservative approach and focus on highly supported phylogenetic relationships mitigates these concerns. Additionally, as *V. cidicii* and *V. navarrensis* become more widely recognized and isolated, increased genomic sampling may reveal additional diversity. Because we have included only isolates characterized as members of the Vulnificus clade (Jiang et al., 2022), it is possible that examination of the Vulnificus clade will need to be repeated as clade definitions are updated and as new isolates are described. For example, *Vibrio floridensis* has been identified as a novel species and is related to *V. vulnificus* (Grant et al., 2023); however, it is not currently included in the Vibrionaceae clades (Jiang et al., 2022).

As *V. cidicii* and *V. navarrensis* become more easily characterized by sequencing applications and our understanding of environmental reservoirs, there will be greater genetic diversity that may alter our studies’ findings. We note an additional 12 *V. cidicii* isolates from the green algae (*Cladophora glomerata*) and seawater (Baltic Sea, Denmark) (Hounmanou et al., 2025) identified after our initial download date in September of 2024. Our findings on virulence are based on genomic content; functional characterization is needed to confirm phenotypic differences. Finally, because isolation of *V. navarrensis* and *V. cidicii* is rare compared to *V. vulnificus* overall, we may not be capturing the full genomic diversity of these two species.

### Conclusions and Future Directions

This comprehensive genomic analysis of 62 genomes reveals that the Vulnificus clade represents a model system for understanding bacterial speciation in aquatic environments. While sharing core metabolic and survival functions, these three species have diverged in key areas: *V. vulnificus* has specialized virulence factors (RTX toxins) enabling severe human pathogenicity, *V. cidicii* has developed metabolic specializations (rhamnose catabolism) for specific environmental niches, and *V. navarrensis* occupies an intermediate position with broad environmental adaptability but reduced virulence potential.

Our results indicate *V. navarrensis* and *V. cidicii* are as closely related as *V. navarrensis* biotype pommerensis is to *V. navarrensis* (95.3% ANI), raising important taxonomic questions that warrant continued monitoring as additional strains become available. Despite the absence of RTX toxins, both *V. navarrensis* and *V. cidicii* continue to be isolated from clinical specimens, suggesting alternative pathogenesis mechanisms that warrant experimental investigation. Future work should include experimental validation of virulence mechanisms and continued genomic surveillance as additional isolates become available to complement these genomic insights.

## ACKNOWLEDGEMENTS

This study did not have any funding. K.A.L: conceptualization, data curation, formal analysis, methodology, project administration, visualization, writing-original draft, writing-reviewing and editing. M.E.J.L.: data curation, formal analysis, methodology, visualization, writing-review and editing.

## REFERENCES

Afgan, E., Baker, D., Batut, B., Van Den Beek, M., Bouvier, D., Čech, M., Chilton, J., Clements, D., Coraor, N., Grüning, B.A. and Guerler, A., 2018. The Galaxy platform for accessible, reproducible and collaborative biomedical analyses: 2018 update. Nucleic acids research, 46(W1), pp.W537–W544.

Alderkamp, A.C., Buma, A.G. and van Rijssel, M., 2007. The carbohydrates of Phaeocystis and their degradation in the microbial food web. Biogeochemistry, 83, pp.99–118.

Ashburner, M., Ball, C.A., Blake, J.A., Botstein, D., Butler, H., Cherry, J.M., Davis, A.P., Dolinski, K., Dwight, S.S., Eppig, J.T. and Harris, M.A., 2000. Gene ontology: tool for the unification of biology. Nature genetics, 25(1), pp.25–29.

Austin, B., 2010. Vibrios as causal agents of zoonoses. Veterinary microbiology, 140(3-4), pp.310–317.

Baker-Austin, C. and Oliver, J.D., 2018. Vibrio vulnificus: new insights into a deadly opportunistic pathogen. Environmental microbiology, 20(2), pp.423–430.

Bier, N., Bechlars, S., Diescher, S., Klein, F., Hauk, G., Duty, O., Strauch, E. and Dieckmann, R., 2013. Genotypic diversity and virulence characteristics of clinical and environmental Vibrio vulnificus isolates from the Baltic Sea region. Applied and environmental microbiology, 79(12), pp.3570–3581.

Centers for Disease Control and Prevention (CDC). 2021. National Vibrio surveillance system: Annual summary, 2019. U.S. Department of Health & Human Services. https://www.cdc.gov/vibrio/php/surveillance/annual-summary-2019.html

Chen L., Yang J., Yu J., Yao Z., Sun L., Shen Y., Jin Q. VFDB: a reference database for bacterial virulence factors. Nucleic Acids Res. 2005; 33:D325–D328.

Chen, H. and Chen, C.Y., 2014. Starvation induces phenotypic diversification and convergent evolution in Vibrio vulnificus. PLoS One, 9(2), p.e88658.

Cuccuru, G., Orsini, M., Pinna, A., Sbardellati, A., Soranzo, N., Travaglione, A., Uva, P., Zanetti, G. and Fotia, G., 2014. Orione, a web-based framework for NGS analysis in microbiology. Bioinformatics, 30(13), pp.1928–1929.

Danin-Poleg, Y., Elgavish, S., Raz, N., Efimov, V. and Kashi, Y., 2013. Genome sequence of the pathogenic bacterium Vibrio vulnificus biotype 3. Genome Announcements, 1(2), pp.10–1128.

de Vries A, Ripley BD (2024). ggdendro: Create Dendrograms and Tree Diagrams Using ‘ggplot2’. R package version 0.2.0, https://andrie.github.io/ggdendro/.

Donati, C., Hiller, N.L., Tettelin, H., Muzzi, A., Croucher, N.J., Angiuoli, S.V., Oggioni, M., Dunning Hotopp, J.C., Hu, F.Z., Riley, D.R. and Covacci, A., 2010. Structure and dynamics of the pan-genome of Streptococcus pneumoniae and closely related species. Genome biology, 11(10), p.R107.

Emms, D.M. and Kelly, S., 2019. OrthoFinder: phylogenetic orthology inference for comparative genomics. Genome biology, 20, pp.1–14.

Farmer, J.J., 1979. Vibrio (“Beneckea”) vulnificus, the bacterium associated with sepsis, septicaemia, and the sea. The Lancet, 314(8148), p.903.

Frey, J., Beck, M., Stucki, U. and Nicolet, J., 1993. Analysis of hemolysin operons in Actinobacillus pleuropneumoniae. Gene, 123(1), pp.51–58.

Gladney LM, Katz LS, Knipe KM, Rowe LA, Conley AB, Rishishwar L, Mariño-Ramírez L, Jordan IK, Tarr CL. Genome sequences of Vibrio navarrensis, a potential human pathogen. Genome announcements. 2014 Dec 24;2(6):e01188–14.

Gladney, L.M. and Tarr, C.L., 2014. Molecular and phenotypic characterization of Vibrio navarrensis isolates associated with human illness. Journal of Clinical Microbiology, 52(11), pp.4070–4074.

Grant, T.A., Jayakumar, J.M., López-Pérez, M. and Almagro-Moreno, S., 2023. Vibrio floridensis sp. nov., a novel species closely related to the human pathogen Vibrio vulnificus isolated from a cyanobacterial bloom. International Journal of Systematic and Evolutionary Microbiology, 73(2), p.005675.

Gurevich, A., Saveliev, V., Vyahhi, N. and Tesler, G., 2013. QUAST: quality assessment tool for genome assemblies. Bioinformatics, 29(8), pp.1072–1075.

Hernández-Cabanyero, C. and Amaro, C., 2020. Phylogeny and life cycle of the zoonotic pathogen Vibrio vulnificus. Environmental Microbiology, 22(10), pp.4133–4148.

Hoang, D. T., Chernomor, O., von Haeseler, A., Minh, B. Q., & Vinh, L. S. (2017). UFBoot2: Improving the Ultrafast Bootstrap Approximation. Molecular Biology and Evolution, 35(2), 518–522. 10.1093/molbev/msx281

Hollis, D.G., Weaver, R.E., Baker, C.N. and Thornsberry, C., 1976. Halophilic Vibrio species isolated from blood cultures. Journal of Clinical Microbiology, 3(4), pp.425–431.

Hounmanou, Y.M.G., Hougbenou, B.G.J., Dougnon, V.T., Hammerl, J.A. and Dalsgaard, A., 2025. Vibrio cidicii genomes recovered from Baltic Sea samples in Denmark. Microbiology Resource Announcements, 14(1), pp.e01121–24.

Huerta-Cepas, J., Forslund, K., Coelho, L.P., Szklarczyk, D., Jensen, L.J., Von Mering, C. and Bork, P., 2017. Fast genome-wide functional annotation through orthology assignment by eggNOG-mapper. Molecular biology and evolution, 34(8), pp.2115–2122.

Huerta-Cepas, J., Szklarczyk, D., Forslund, K., Cook, H., Heller, D., Walter, M.C., Rattei, T., Mende, D.R., Sunagawa, S., Kuhn, M. and Jensen, L.J., 2016. eggNOG 4.5: a hierarchical orthology framework with improved functional annotations for eukaryotic, prokaryotic and viral sequences. Nucleic acids research, 44(D1), pp.D286–D293.

Hunt, D.E., David, L.A., Gevers, D., Preheim, S.P., Alm, E.J. and Polz, M.F., 2008. Resource partitioning and sympatric differentiation among closely related bacterioplankton. Science, 320(5879), pp.1081–1085.

Jain, C., Rodriguez-R, L. M., Phillippy, A. M., Konstantinidis, K. T., & Aluru, S. (2018). High throughput ANI analysis of 90K prokaryotic genomes reveals clear species boundaries. Nature Communications, 9(1). 10.1038/s41467-018-07641-9

Janda, J.M., 2020. Clinical decisions: detecting vibriosis in the modern era. Clinical Microbiology Newsletter, 42(6), pp.45–50.

Jia, B., Raphenya, A.R., Alcock, B., Waglechner, N., Guo, P., Tsang, K.K., Lago, B.A., Dave, B.M., Pereira, S., Sharma, A.N. and Doshi, S., 2016. CARD 2017: expansion and model-centric curation of the comprehensive antibiotic resistance database. Nucleic acids research, p.gkw1004.

Jiang, C., Tanaka, M., Nishikawa, S., Mino, S., Romalde, J.L., Thompson, F.L., Gomez-Gil, B. and Sawabe, T., 2022. Vibrio Clade 3.0: new Vibrionaceae evolutionary units using genome-based approach. Current microbiology, 79, pp.1–15.

Jones, M.K. and Oliver, J.D., 2009. Vibrio vulnificus: disease and pathogenesis. Infection and immunity, 77(5), pp.1723–1733.

Jores J, Appel B, Lewin A. Vibrio navarrensis biotype pommerensis: a new biotype of V. navarrensis isolated in the German Baltic Sea. Systematic and applied microbiology. 2007 Jan 19;30(1):27–30.

Kalyaanamoorthy, S., Minh, B. Q., Wong, T. K. F., von Haeseler, A., & Jermiin, L. S. (2017). ModelFinder: fast model selection for accurate phylogenetic estimates. Nature Methods, 14(6), 587–589. 10.1038/nmeth.4285

Kanehisa, M., Sato, Y., and Morishima, K.; BlastKOALA and GhostKOALA: KEGG tools for functional characterization of genome and metagenome sequences. J. Mol. Biol. 428, 726–731 (2016).

Kannan, S., Sharma, S., Ciufo, S., Clark, K., Turner, S., Kitts, P.A., Schoch, C.L., DiCuccio, M. and Kimchi, A., 2023. Collection and curation of prokaryotic genome assemblies from type strains at NCBI. International Journal of Systematic and Evolutionary Microbiology, 73(1), p.005707.

Kaspar, C.A. and Tamplin, M.L., 1993. Effects of temperature and salinity on the survival of Vibrio vulnificus in seawater and shellfish. Applied and environmental microbiology, 59(8), pp.2425–2429.

Keck, F., 2020. Handling biological sequences in R with the bioseq package. Methods Ecol Evol. 11: 1728–1732 [online]

Koton, Y., Gordon, M., Chalifa-Caspi, V. and Bisharat, N., 2015. Comparative genomic analysis of clinical and environmental Vibrio vulnificus isolates revealed biotype 3 evolutionary relationships. Frontiers in Microbiology, 5, p.803.

Lai, K.P., Tsang, C.F., Li, L., Yu, R.M.K. and Kong, R.Y.C., 2022. Microplastics act as a carrier for wastewater-borne pathogenic bacteria in sewage. Chemosphere, 301, p.134692.

Letunic and Bork (2024) Nucleic Acids Res doi: 10.1093/nar/gkae268

Letunic, I. and Bork, P., 2024. Interactive Tree of Life (iTOL) v6: recent updates to the phylogenetic tree display and annotation tool. Nucleic Acids Research, p.gkae268.

Lo, L.S.H., Tong, R.M.K., Chan, W., Ho, W. and Cheng, J., 2025. Bacterial pathogen assemblages on microplastic biofilms in coastal waters. Marine Pollution Bulletin, 216, p.117958.

López-Pérez, M., Jayakumar, J.M., Haro-Moreno, J.M., Zaragoza-Solas, A., Reddi, G., Rodriguez-Valera, F., Shapiro, O.H., Alam, M. and Almagro-Moreno, S., 2019. Evolutionary model of cluster divergence of the emergent marine pathogen Vibrio vulnificus: from genotype to ecotype. MBio, 10(1), pp.10–1128.

Lydon KA, Robertson MJ, Lipp EK. Patterns of triclosan resistance in Vibrionaceae. PeerJ. 2018 Jul 12;6:e5170.

Lydon, K.A., Grim, C., Robertson, M.J. and Lipp, E.K., 2024. Draft genome sequence of multidrug-resistant Vibrio navarrensis strain DA9 isolated from a coastal canal in the Florida Keys (USA). Microbiology Resource Announcements, 13(1), pp.e00789–23.

Macián MC, Arias CR, Aznar R, Garay E, Pujalte MJ. Identification of Vibrio spp.(other than V. vulnificus) recovered on CPC agar from marine natural samples. International Microbiology. 2000 Mar 1;3(1):51–3.

Mikheenko, A., Prjibelski, A., Saveliev, V., Antipov, D. and Gurevich, A., 2018. Versatile genome assembly evaluation with QUAST-LG. Bioinformatics, 34(13), pp.i142–i150.

Mikheenko, A., Valin, G., Prjibelski, A., Saveliev, V. and Gurevich, A., 2016. Icarus: visualizer for de novo assembly evaluation. Bioinformatics, 32(21), pp.3321–3323.

Minh, B.Q., Schmidt, H.A., Chernomor, O., Schrempf, D., Woodhams, M.D., Von Haeseler, A. and Lanfear, R., 2020. IQ-TREE 2: new models and efficient methods for phylogenetic inference in the genomic era. Molecular biology and evolution, 37(5), pp.1530–1534.

Morrison, S.S., Pyzh, R., Jeon, M.S., Amaro, C., Roig, F.J., Baker-Austin, C., Oliver, J.D. and Gibas, C.J., 2014, November. Impact of analytic provenance in genome analysis. In BMC genomics (Vol. 15, pp.1–11). BioMed Central.

Mühlenbruch, M., Grossart, H.P., Eigemann, F. and Voss, M., 2018. Mini-review: Phytoplankton-derived polysaccharides in the marine environment and their interactions with heterotrophic bacteria. Environmental microbiology, 20(8), pp.2671–2685.

NCBI https://www.ncbi.nlm.nih.gov/nuccore/JBEWCK000000000.1 accessed 9/21/2024

Nilsson, W.B., Paranjype, R.N., DePaola, A. and Strom, M.S., 2003. Sequence polymorphism of the 16S rRNA gene of Vibrio vulnificus is a possible indicator of strain virulence. Journal of clinical microbiology, 41(1), pp.442–446.

Oliver, J.D., 2005. Wound infections caused by Vibrio vulnificus and other marine bacteria. Epidemiology & Infection, 133(3), pp.383–391.

Oliver, J.D., Wear, J.E., Thomas, M.B., Warner, M. and Linder, K., 1986. Production of extracellular enzymes and cytotoxicity by Vibrio vulnificus. Diagnostic microbiology and infectious disease, 5(2), pp.99–111.

Orata FD, Xu Y, Gladney LM, Rishishwar L, Case RJ, Boucher Y, Jordan IK, Tarr CL. Characterization of clinical and environmental isolates of Vibrio cidicii sp. nov., a close relative of Vibrio navarrensis. International Journal of Systematic and Evolutionary Microbiology. 2016 Oct 1;66(10):4148–55.

Page, A.J., Cummins, C.A., Hunt, M., Wong, V.K., Reuter, S., Holden, M.T., Fookes, M., Falush, D., Keane, J.A. and Parkhill, J., 2015. Roary: rapid large-scale prokaryote pan genome analysis. Bioinformatics, 31(22), pp.3691–3693.

R Core Team (2021). R: A language and environment for statistical computing. R Foundation for Statistical Computing, Vienna, Austria.

Rodriguez-R, L.M., Conrad, R.E., Viver, T., Feistel, D.J., Lindner, B.G., Venter, S.N., Orellana, L.H., Amann, R., Rossello-Mora, R. and Konstantinidis, K.T., 2024. An ANI gap within bacterial species that advances the definitions of intra-species units. MBio, 15(1), pp.e02696–23.

Roig, F.J., González-Candelas, F., Sanjuán, E., Fouz, B., Feil, E.J., Llorens, C., Baker-Austin, C., Oliver, J.D., Danin-Poleg, Y., Gibas, C.J. and Kashi, Y., 2018. Phylogeny of Vibrio vulnificus from the analysis of the core-genome: implications for intra-species taxonomy. Frontiers in microbiology, 8, p.2613.

Sawabe, T., Kita-Tsukamoto, K. and Thompson, F.L., 2007. Inferring the evolutionary history of vibrios by means of multilocus sequence analysis. Journal of bacteriology, 189(21), pp.7932–7936.

Sawabe, T., Ogura, Y., Matsumura, Y., Feng, G., Amin, A.R., Mino, S., Nakagawa, S., Sawabe, T., Kumar, R., Fukui, Y. and Satomi, M., 2013. Updating the Vibrio clades defined by multilocus sequence phylogeny: proposal of eight new clades, and the description of Vibrio tritonius sp. nov. Frontiers in microbiology, 4, p.414.

Schwartz K, Borowiak M, Deneke C, Balau V, Metelmann C, Strauch E. Complete and Circularized Genome Assembly of a Human Isolate of Vibrio navarrensis Biotype pommerensis with MiSeq and MinION Sequence Data. Microbiology Resource Announcements. 2021 Feb 4;10(5):e01435–20.

Schwartz K, Kukuc C, Bier N, Taureck K, Hammerl JA, Strauch E. Diversity of Vibrio navarrensis revealed by genomic comparison: veterinary isolates are related to strains associated with human illness and sewage isolates while seawater strains are more distant. Frontiers in microbiology. 2017 Sep x6;8:1717.

Seemann, T. (2016). ABRicate: mass screening of contigs for antiobiotic resistance genes. https://github.com/tseemann/abricate

Seemann, T., 2014. Prokka: rapid prokaryotic genome annotation. Bioinformatics, 30(14), pp.2068–2069.

Smet, A., Yahara, K., Rossi, M., Tay, A., Backert, S., Armin, E., Fox, J.G., Flahou, B., Ducatelle, R., Haesebrouck, F. and Corander, J., 2018. Macroevolution of gastric Helicobacter species unveils interspecies admixture and time of divergence. The ISME Journal, 12(10), pp.2518–2531.

Tang, A. D., Soulette, C. M., van Baren, M. J., Hart, K., Hrabeta-Robinson, E., Wu, C. J., & Brooks, A. N. (2020). Full-length transcript characterization of SF3B1 mutation in chronic lymphocytic leukemia reveals downregulation of retained introns. Nature Communications, 11(1). 10.1038/s41467-020-15171-6

The Galaxy Community. The Galaxy platform for accessible, reproducible, and collaborative data analyses: 2024 update, Nucleic Acids Research, 2024;, gkae410, 10.1093/nar/gkae410

Urdaci MC, Marchand M, Ageron E, Arcos JM, Sesma B, Grimont PA. Vibrio navarrensis sp. nov., a species from sewage. International Journal of Systematic and Evolutionary Microbiology. 1991 Apr 1;41(2):290–4.

Wan, S.H., Xu, Y., Xu, W., Leung, S.K., Yu, E.Y. and Yung, C.C., 2025. Environmental Heterogeneity Drives Ecological Differentiation in Vibrio Populations Across Subtropical Marine Habitats. Environmental Microbiology, 27(5), p.e70107.

